# Optimisation and Validation of a UHPLC-MS/MS Assay Using Timed Post-Column Ammonium Fluoride Infusion for Sensitive Serum Steroids Quantitation

**DOI:** 10.64898/2026.10.27.762247

**Authors:** Věra Dosedělová, James M. Hawley, Eka Melson, Lina Schiffer, Karl-Heinz Storbeck, Wiebke Arlt

## Abstract

Steroid metabolomics has potential for identifying biomarkers and improving disease diagnosis. However, poor ionisation efficiency and low circulating concentrations of some steroids limit the broader clinical applications. Here, we developed a ultra-high-performance liquid chromatography – tandem mass spectrometry (UHPLC-MS/MS) assay for quantitation of 18 steroids in serum encompassing the mineralocorticoid, glucocorticoid and androgen biosynthesis pathways, including both classic and 11-oxygenated androgens. Steroid extraction from 100 μL of serum was performed using supported liquid extraction with ethyl acetate elution. Ionisation efficiency of steroids was investigated using formic acid in the mobile phase and post-column infusion of ammonium fluoride. Ammonium fluoride enhanced the signal intensity of 3-keto-Δ4 steroids up to ninefold, however A-ring reduced steroids exhibited greater ionisation with formic acid. Thus, to optimise sensitivity, a timed post-column infusion of 2 mmol/L ammonium fluoride was employed during chromatographic separation. The lower limits of quantitation ranged from 0.04 to 2.1 nmol/L. Accuracy ranged from 82 to 113% and intra- and inter-assay imprecision was <10% and <13%, respectively. The assay was applied to analyse morning serum steroid concentrations in healthy individuals (55 women, 44 men; age-matched). Serum progesterone, 11-deoxycorticosterone, dehydroepiandrosterone and androstenedione were significantly higher in women; in contrast, testosterone and 5α-dihydrotestosterone were higher in men. In summary, we have successfully developed and validated a UHPLC-MS/MS assay for multi-steroid profiling ad applied it to healthy individuals of both sexes.

## 1 Introduction

Steroid hormones are essential for regulating a multitude of physiological processes, including reproduction, development, and metabolism. Steroids are primarily biosynthesised from cholesterol in the adrenal glands and gonads [1]. The combination of *de novo* steroid hormone biosynthesis and subsequent peripheral metabolism generates a complex circulating steroid metabolome that is altered in various pathological conditions. Consequently, steroid metabolome profiling has considerable potential for elucidating disease mechanisms, identifying biomarkers, and improving disease diagnosis and monitoring. Examples of disorders characterised by distinct alterations in the circulating steroid metabolome include adrenocortical carcinoma [2], congenital adrenal hyperplasia [3], polyendocrine metabolic ovarian syndrome (PMOS) [4,5], and Cushing’s syndrome [3].

Immunoassays remain widely used for steroid analysis especially in clinical laboratories because of their cost-effectiveness, simplicity, and short turn-around times. However, these methods often lack analytical specificity and may overestimate steroid concentrations, particularly at low levels due to cross-reactivity with structurally similar compounds [6–8]. Conversely, liquid chromatography – tandem mass spectrometry (LC-MS/MS) offers superior sensitivity, specificity, and multiplexing capability. Multiplexed steroid analyses support a more comprehensive characterisation of the metabolome enabling a holistic assessment of the relationships between different steroidogenic pathways. Furthermore, LC-MS/MS steroid profiling combined with machine learning has emerged as a promising strategy for improving diagnostic accuracy, patient stratification, and disease classification [2,9,10].

Despite these advantages, comprehensive steroid profiling remains analytically challenging because many steroids occur at low circulating concentrations, exhibit poor ionisation efficiency, and exist as structurally similar isomers and stereoisomers that require adequate chromatographic separation. Mobile phase additives such as ammonium fluoride (NH_4_F), formic acid (FA) and ammonium acetate are commonly employed to improve ionisation efficiency and sensitivity of steroid detection [8,11,12]. In addition, oxime derivatisation has been used to enhance steroid ionisation and specificity [13,14]. However, derivatisation increases sample preparation complexity and requires additional method optimisation. Sample preparation for serum steroid analysis by LC-MS/MS typically involves protein precipitation followed by liquid–liquid extraction (LLE) or solid-phase extraction (SPE), with some methods requiring up to 500 μL serum [8,15–18].

Herein, we present an optimised supported liquid extraction (SLE) protocol for the extraction of 18 serum steroids using a low sample volume followed by UHPLC-MS/MS quantification combining FA in the mobile phase and a timed post-column infusion of NH_4_F to selectively enhance ionisation according to steroid A-ring structure. The method was subsequently applied to assess steroid concentrations in healthy individuals.

## 2 Material and Methods

### 2.1 Chemicals

The reference steroid standards and internal standards (IS) used in the assay are summarised in Supplementary Table S1. Certified reference materials (CRM) of concentration 1 mg/mL or 100 μg/mL in methanol/acetonitrile were used as stock solutions for calibrators, quality control (QC) samples and IS. CRMs were obtained from Cerilliant (Toronto Research Chemicals, TX, USA), Lipomed (Arlesheim, Switzerland) and Cayman Chemical (Ann Arbor, MI, USA). Where CRMs were unavailable, 1 mg/mL stock solutions were gravimetrically prepared in methanol from powder (purity > 98%) and stored at −20°C. Separate stock solutions of different lot numbers were used for preparation of calibrators and QCs. A combined IS solution (10-250 ng/mL) was prepared in methanol and stored at −80°C. LC-MS grade methanol, water, ethyl acetate (EtOAc) and hexane were purchased from Honeywell (CHROMASOLV^TM^, Seelze, Germany). LC-MS grade FA (99%) was obtained from Biosolve (Dieuze, France). Phosphate buffered saline (PBS), bovine serum albumin (BSA), NH_4_F, and HPLC grade dichloromethane (DCM) were obtained from Merck (Gillingham, UK). LC-MS grade methyl tert-butyl ether (MTBE) was purchased from Rathburn Chemicals (Walkerburn, UK).

### 2.2 Preparation of Calibrators and QC Samples

Intermediate solutions for calibrators and QCs were prepared in methanol from reference standard stock solutions. The concentration of intermediate solution was 600 ng/mL for all steroids, except for cortisol (10,000 ng/mL) and cortisone (2,000 ng/mL). Calibrators (Supplementary Table S2) and QC samples (Supplementary Table S3) were prepared gravimetrically from intermediate solutions using PBS 2% BSA which served as a surrogate matrix for human serum. Calibrators (n = 9) ranged from 0 to 30 ng/mL (87-105 nmol/L), except for cortisol, i.e., 0-500 ng/mL (1,380 nmol/L), and cortisone, i.e., 0-100 ng/mL (277 nmol/L). QC samples (n = 7) ranged from 0.07 (0.2-0.3) to 24 ng/mL (69-84 nmol/L), except for cortisol, i.e., 1.2 (3.4) - 400 ng/mL (1,100 nmol/L), and cortisone, i.e., 0.25 (0.7) - 80 ng/mL (222 nmol/L). Aliquots of calibrators and QCs were stored at −80°C.

### 2.3 Sample Preparation

Calibrators, QC and serum samples (100 μL) were pipetted into the wells of a 96-well plate (2 mL; Porvair Sciences Ltd, Wrexham, UK) followed by the addition of 20 μL IS solution and 100 μL LC-MS grade water (CHROMASOLV^TM^) to each well. The plate was heat-sealed, vortex mixed for 5 min, centrifuged (10 min, 1,900 g) and transferred to a 200 µL SLE plate (ISOLUTE® SLE+, Biotage, Uppsala, Sweden). A positive pressure of 3 psi was applied for 30 s (PRESSURE+ 96, Biotage) to load the sample onto the diatomaceous earth, the SLE plates were equilibrated for 5 min at room temperature. Steroids were eluted with 1 mL of EtOAc by gravity into a 2 mL 96-well plate (Porvair Sciences) followed by the application of a positive pressure of 5 psi for 10 s. SLE eluates were subsequently dried under nitrogen (gas flow: 50 L/min, gas temperature: 25°C, plate temperature: 25°C) using a TurboVap® 96 Dual evaporation system (Biotage). Prior to UHPLC-MS/MS analysis, samples were reconstituted in 100 µL of 50% methanol/water (v/v), vortex mixed for 10 min and centrifuged (10 min, 1,900 g).

### 2.4 UHPLC-MS/MS Analysis

UHPLC-MS/MS analysis was performed on an ACQUITY Premier UHPLC system coupled to a triple quadrupole mass spectrometer Xevo TQ-XS with a Z-spray ion source (Waters, Wilmslow, UK). MassLynx software V4.2 was used for instrument control and data acquisition. TargetLynx XS V4.2 was used for data processing. A Luna Omega Polar C18 column (1.6 µm, 100Å, 100 x 2.1 mm; Phenomenex, Macclesfield, UK) was combined with an ACQUITY column in-line filter (Waters). The column temperature was maintained at 60°C. The mobile phases consisted of (A) water with 0.01% FA and (B) methanol with 0.01% FA. The injection volume was 10 μL. Steroids were separated at a flow rate of 0.4 mL/min using a gradient elution: 0-0.5 min, 50% B; 0.5-8.4 min, 50-65% B; 8.4-9.6 min, 95% B (wash step); 9.6-11.5 min, 50% B (equilibration step). A timed post-column infusion of 2 mmol/L NH_4_F in 50% methanol/water (v/v) was employed to enhance detection sensitivity. The post-column infusion was delivered at a flow rate of 5 μL/min between 1.1 and 5.15 min of the analytical run. An electrospray ionisation ion source was operated in the positive ionisation mode. The capillary voltage was set to 1.5 kV, while the desolvation temperature and gas flow were maintained at 600°C and 1200 L/hr, respectively. The cone and nebuliser gas flow were 150 L/hr and 7 bar, respectively. The source temperature was 150 °C. Steroids were detected in the multiple reaction monitoring mode. All mass transitions with their respective cone voltages, collision energies and retention times are summarised in Table 1.

**Table 1.** Multiple reaction monitoring transitions for MS/MS detection (CV – cone voltage, CE – collision energy, t_R_ – retention time).

| Abbr. | Steroid name<br>Internal standard | Quantifier / Qualifier / Internal standard |  |  |  |  |
| --- | --- | --- | --- | --- | --- | --- |
|  |  | Precursor ion | Mass transition | CV (V) | CE (eV) | t<sub>R</sub> (min) |
| **Prog** | Progesterone<br>Prog-<sup>13</sup>C3 | [M+H]<sup>+</sup> | 315.2 > 97.0 | 25 | 20 | 7.97 |
|  |  |  | 315.2 > 109.0 | 25 | 24 |  |
|  |  |  | 318.2 > 100.0 | 25 | 20 |  |
| **17OHP** | 17α-Hydroxyprogesterone<br>17OHP-<sup>13</sup>C3 | [M+H]<sup>+</sup> | 331.2 > 97.0 | 25 | 25 | 5.39 |
|  |  |  | 331.2 > 109.0 | 25 | 28 |  |
|  |  |  | 334.2 > 100.0 | 25 | 25 |  |
| **DOC** | 11-Deoxycorticosterone<br>DOC-<sup>13</sup>C3 | [M+H]<sup>+</sup> | 331.2 > 97.0 | 25 | 22 | 4.86 |
|  |  |  | 331.2 > 109.0 | 25 | 27 |  |
|  |  |  | 334.2 > 100.0 | 25 | 22 |  |
| **B** | Corticosterone<br>B-<sup>13</sup>C4 | [M+H]<sup>+</sup> | 347.2 > 121.0 | 20 | 23 | 3.17 |
|  |  |  | 347.2 > 97.0 | 20 | 25 |  |
|  |  |  | 351.3 > 123.1 | 20 | 23 |  |
| **S** | 11-Deoxycortisol<br>S-<sup>13</sup>C3 | [M+H]<sup>+</sup> | 347.2 > 97.0 | 25 | 23 | 3.39 |
|  |  |  | 347.2 > 109.0 | 25 | 26 |  |
|  |  |  | 350.2 > 100.0 | 25 | 23 |  |
| **F** | Cortisol<br>F-d4 | [M+H]<sup>+</sup> | 363.2 > 97.0 | 25 | 25 | 2.16 |
|  |  |  | 363.2 > 121.0 | 25 | 25 |  |
|  |  |  | 367.2 > 97.0 | 25 | 25 |  |
| **E** | Cortisone<br>E-<sup>13</sup>C3 | [M+H]<sup>+</sup> | 361.2 > 163.1 | 27 | 23 | 1.94 |
|  |  |  | 361.2 > 121.0 | 27 | 28 |  |
|  |  |  | 364.2 > 166.1 | 27 | 23 |  |
| **DHEA** | Dehydroepiandrosterone<br>DHEA-d5 | [M+H-H<sub>2</sub>O]<sup>+</sup> | 271.2 > 213.1 | 12 | 16 | 5.30 |
|  |  |  | 271.2 > 253.1 | 12 | 12 |  |
|  |  |  | 276.2 > 218.1 | 12 | 16 |  |
| **A4** | Androstenedione<br>A4-<sup>13</sup>C3 | [M+H]<sup>+</sup> | 287.2 > 97.0 | 23 | 18 | 4.43 |
|  |  |  | 287.2 > 109.0 | 23 | 22 |  |
|  |  |  | 290.2 > 100.0 | 23 | 18 |  |
| **5α-dione** | 5α-Androstane-3,17-dione<br>5α-dione-<sup>13</sup>C3 | [M+H]<sup>+</sup> | 289.2 > 253.1 | 25 | 15 | 5.95 |
|  |  |  | 289.2 > 213.1 | 25 | 18 |  |
|  |  |  | 292.2 > 256.1 | 25 | 15 |  |
| **An** | 5α-Androsterone<br>An-d4 | [M+H]<sup>+</sup> | 291.2 > 255.1 | 20 | 14 | 7.84 |
|  |  |  | 291.2 > 273.1 | 20 | 8 |  |
|  |  |  | 295.2 > 259.1 | 20 | 14 |  |
| **T** | Testosterone<br>T-<sup>13</sup>C3 | [M+H]<sup>+</sup> | 289.2 > 97.0 | 25 | 20 | 5.03 |
|  |  |  | 289.2 > 109.0 | 25 | 24 |  |
|  |  |  | 292.2 > 100.0 | 25 | 20 |  |
| **DHT** | 5α-Dihydrotestosterone<br>DHT-d3 | [M+H]<sup>+</sup> | 291.2 > 255.2 | 25 | 14 | 6.44 |
|  |  |  | 291.2 > 159.1 | 25 | 20 |  |
|  |  |  | 294.2 > 258.2 | 25 | 14 |  |
| **3α-adiol** | 5α-Androstane-3α,17β-diol<br>3α-adiol-d3 | [M+H-2H<sub>2</sub>O]<sup>+</sup> | 257.2 > 147.0 | 40 | 18 | 7.47 |
|  |  |  | 257.2 > 135.0 | 20 | 17 |  |
|  |  |  | 260.2 > 147.0 | 40 | 18 |  |
| **11OHA4** | 11β-Hydroxyandrostenedione<br>11OHA4-<sup>13</sup>C3 | [M+H]<sup>+</sup> | 303.2 > 267.1 | 20 | 15 | 2.77 |
|  |  |  | 303.2 > 285.2 | 20 | 14 |  |
|  |  |  | 306.2 > 270.2 | 20 | 15 |  |
| **11KA4** | 11-Ketoandrostenedione<br>11KA4-<sup>13</sup>C3 | [M+H]<sup>+</sup> | 301.2 > 257.1 | 25 | 23 | 2.17 |
|  |  |  | 301.2 > 121.0 | 25 | 25 |  |
|  |  |  | 304.2 > 260.1 | 25 | 23 |  |
| **11KT** | 11-Ketotestosterone<br>11KT-<sup>13</sup>C3 | [M+H]<sup>+</sup> | 303.2 > 121.0 | 25 | 25 | 2.48 |
|  |  |  | 303.2 > 259.1 | 25 | 22 |  |
|  |  |  | 306.2 > 124.1 | 25 | 25 |  |
| **11OHT** | 11β-Hydroxytestosterone<br>11OHT-<sup>13</sup>C3 | [M+H]<sup>+</sup> | 305.2 > 269.2 | 25 | 14 | 3.07 |
|  |  |  | 305.2 > 147.0 | 25 | 23 |  |
|  |  |  | 308.2 > 272.2 | 25 | 14 |  |

### 2.5 Validation

The UHPLC-MS/MS assay was validated according to the Bioanalytical Method Validation and Study Sample Analysis Guideline [19] and Bioanalytical Method Validation Guidance for Industry [20].

#### 2.5.1 Specificity

To evaluate specificity, the separation of common steroid isobars of the same *m/z* was investigated. Steroids were either spiked into PBS 2% BSA at a final concentration of 20 ng/mL, or commercially available quality controls were analysed (MassCheck® Steroid Panel 1 Serum Controls; Chromsystems Instruments & Chemicals GmbH, Munich, Germany). The following isobars were investigated: epitestosterone for testosterone, aldosterone and prednisolone for cortisone, 21-deoxycortisol for 11-deoxycortisol, and 20α/20β-dihydrocortisone for cortisol.

#### 2.5.2 Calibration curves and ranges

Calibration curves were prepared in PBS 2% BSA as described in Section 2.2 in the ranges 0.257-500 ng/mL (0.7-1379 nmol/L) for cortisol, 0.051-100 ng/mL (0.1-277 nmol/L) for cortisone and 0.015-30 ng/mL (∼0.04-105 nmol/L) for the remaining steroids. For each analyte, the ratio of analyte peak area to IS peak area was plotted against the nominal concentration. A 1/X weighted linear regression fit was considered acceptable if a coefficient of determination (R^2^) ≥ 0.99 and accuracy of the back-calculated concentrations was within ±20% of the nominal concentration of the lower limit of quantitation (LLOQ) and ±15% for all other concentrations, including the upper limit of quantitation (ULOQ). LLOQs were determined as the lowest quantifiable calibration level with acceptable accuracy ±20% of the nominal concentration.

#### 2.5.3 Carry-over

Carry-over was determined by sequential injections of Cal 9 prepared in 50% methanol/water (v/v) followed by injection of the blank sample (50% methanol/water, v/v). Carry-over in the blank samples was considered acceptable if the analyte response was less than 20% of the response at the LLOQ and 5% of the response for the IS.

#### 2.5.4 Dilution Integrity

The QC sample was freshly prepared in PBS 2% BSA at a concentration 2-fold higher than the ULOQ (2xULOQ QC). The sample was then diluted 10-fold and 40-fold using PBS 2% BSA and analysed in 5 replicates. The measured concentration was back-calculated according to the corresponding dilution factor and compared with the nominal value. Accuracy within ±15% of the nominal concentration and imprecision (expressed as coefficient of variation, CV) below 15% were considered acceptable.

#### 2.5.5 Accuracy and Imprecision

Seven levels of QC samples were prepared in PBS 2% BSA (Supplementary Table S3) to account for the different circulating concentrations and detection sensitivities of each steroid. Accuracy and imprecision were evaluated at all seven levels, but for simplicity only four concentration levels, i.e., QC at LLOQ, low QC, medium QC, and high QC are reported for each steroid.

Intra-assay accuracy and imprecision were determined by analysing five replicates of each QC level within one batch. Inter-assay accuracy and imprecision were determined by analysing QC samples over four separate batches. In addition, imprecision was evaluated in triplicate using a pooled serum sample spiked at four concentration levels (intra-assay CV) and analysed over four different batches (inter-assay CV). Accuracy within ±15% of the nominal concentration and within ±20% at the LLOQ was considered acceptable. Imprecision below 15% and below 20% at the LLOQ was considered acceptable.

#### 2.5.6 Absolute Recovery

To determine absolute recovery, serum samples and PBS 2% BSA were spiked with spiking solutions prepared in 50% methanol/water (v/v) at three QC levels (low = QC 3, medium = QC 4, high = QC 6; Supplementary Table S3) and three replicates. Recovery was calculated as follows: (measured concentration – endogenous concentration) / target concentration × 100. Recovery between 80% and 120% was deemed acceptable.

#### 2.5.7 Matrix Effects

Matrix effects were assessed by post-extraction spiking of analytes and IS into extracted matrix samples / PBS 2% BSA and comparing the resulting responses to that of their corresponding solvent standards prepared in 50% methanol/water (v/v). After SLE elution and solvent evaporation, serum samples from six donors were spiked in triplicate at low (= QC 3) and high QC (= QC 6) levels and IS. For each serum sample, endogenous steroid levels were determined as well. In addition, naturally haemolysed and lipaemic serum samples were extracted and spiked with only IS to determine matrix effects. Matrix factors were calculated as follows: (post-extraction peak area of spike – peak area of endogenous levels) / peak area of solvent standard. Subsequently, IS-normalised matrix effect was determined as a ratio of analyte and IS matrix factors. Matrix effects between 80% and 120% were considered acceptable.

#### 2.5.8 Stability

The stability was investigated in serum samples collected from four women and four men, and a spiked pooled serum sample, all analysed in triplicate. The serum samples were analysed on the day of blood collection (= time zero). The time zero plate was resealed, stored at −20°C for 72 hours, and reinjected to investigate freezer-storage stability. To assess freeze-thaw stability, 450 μL aliquots of each sample were stored at −80°C and subjected to three freeze-thaw cycles, with a minimum of 24 hours between each cycle. Autosampler stability at 10°C was determined by analysing samples on the day of preparation and reinjecting them after 72 hours of storage in the autosampler. The mean concentration bias within ±15% of the nominal (time zero) concentration was considered acceptable.

### 2.6 Collection of Serum Samples from Healthy Volunteers

The collection of serum from healthy volunteers was approved by the Science, Technology, Engineering and Mathematics Ethical Review Committee of the University of Birmingham, Birmingham, UK (ERN_17-0494, ERN_17-0494B) and Imperial College Research Ethics Committee, London, UK (Ref. 7561939). Informed, written consent was obtained from all individuals included in this study. Exclusion criteria were any acute or chronic disease affecting steroid biosynthesis or metabolism and the intake of any medication known to interfere with steroid biosynthesis or metabolism including hormonal contraceptives. A standardised questionnaire was used to record demographic data.

Serum samples were collected in the morning before 11 a.m. from 55 premenopausal female and 44 male healthy age-matched individuals (women median age 37 (range 22-55) years; men median age 38 (range 22-60) years). Blood was collected into BD Vacutainer™ Serum Tubes (Becton Dickinson, Plymouth, UK). After blood coagulation (approx. 45 min, room temperature), samples were centrifuged (10 min, 3,000 g), serum was aliquoted and stored at −80°C until UHPLC-MS/MS analysis.

### 2.7 Data Analysis

Bar plots showing mean and SD were created using GraphPad Prism (version 10.2.3; GraphPad Software, Boston, MA, USA). Boxplots show the median and the interquartile range (lower and upper quartile boundaries). Individual data points are shown as overlaid dots. To compare steroid concentrations between males and females, data distribution was first assessed using histograms, Q-Q plots and Shapiro-Wilk test. In case of normal distribution, steroid concentrations were compared using a two-sample t-test. Otherwise, the Mann-Whitney U test was applied. P-values < 0.05 were considered statistically significant. Data visualisation and statistical analyses were performed using R (version 4.4.0; R Foundation for Statistical Computing, Vienna, Austria), RStudio (version 2026.07.1; Posit Software, PBC, Boston, MA, USA), and packages ggplot2 (version 4.0.3), patchwork (version 1.3.2), scales (version 1.4.0), tidyverse (version 2.0.0), rstatix (version 1.1.0), and dplyr (version 1.2.1). ChatGPT (OpenAI, San Francisco, CA, USA) was used to assist with the development and troubleshooting of R code for statistical analysis and data visualisation. All analyses were performed and verified by authors.

## 3 Results

### 3.1 Chromatographic Separation

The main aspects of the UHPLC-MS/MS method development and optimisation were chromatographic separation and ionisation efficiency. Chromatographic separation of the target steroids was compared using 50 mm and 100 mm Luna Omega Polar C18 columns. The initial mobile phase consisted of 0.1% FA in both the water and methanol phases. The main focus was to achieve baseline separation of structural isomers, i.e., corticosterone and 11-deoxycortisol (*m/z* 347), 11OHA4 and 11KT (*m/z* 303), and DOC and 17OHP (*m/z* 331). Under our conditions, we were unable to achieve separation of these structural isomers on the 50 mm column in a suitable timeframe (Supplementary Figure S1A). However, complete baseline separation of the structural isomers was achieved, with an analysis time of 11.5 min on the 100 mm column (Figure S1B). The higher resolving properties were also beneficial for the implementation of the timed NH_4_F post-column infusion as presented in Section 3.2.

### 3.2 Steroid Ionisation Efficiency

Steroid ionisation efficiency was investigated by comparing the signal response of a 2 ng/mL standard solution in 50% methanol/water (v/v) using FA in mobile phases and post-column infusion of NH_4_F. The ionisation efficiency was evaluated for both FA and NH_4_F separately (Figure 1A) and in combination (Figure 1B). Ammonium fluoride enhanced the signal intensity of 3-keto-Δ4 steroids (i.e., Prog, 17OHP, DOC, B, S, F, E, A4, T, 11OHA4, 11KA4, 11KT, 11OHT) by four- to ninefold compared to FA alone (Figure 1A). When FA was present in mobile phases, NH_4_F enhanced the signal intensity of 3-keto-Δ4 steroids by two- to fivefold (Figure 1B). The signal of DHT (5α-reduced steroid) was enhanced twofold during NH_4_F infusion (Figure 1A). Conversely, 5α-dione (5α-reduced steroid), DHEA (3b-hydroxy steroid), An and 3α-adiol (3a-hydroxy steroids) ionised less well using the NH_4_F infusion relative to FA alone. For instance, 3α-adiol was undetectable using the NH_4_F infusion. Since 3-hydroxy and 5α-reduced steroids ionised poorly regardless of the additive, a timed post-column infusion of NH_4_F was utilised during chromatographic separation (1.1 min – 5.15 min; Figure 2) to enhance Δ4 steroid ionisation, without suppressing the ionisation of poorly ionising steroids DHEA (t_R_ = 5.3 min), 5α-dione (t_R_ = 5.95 min), 3α-adiol (t_R_ = 7.47 min) and An (t_R_ =7.84). Using this strategy, it was not possible to introduce infusion for 17OHP (t_R_ = 5.39 min), DHT (t_R_ = 6.44 min), and Prog (t_R_ = 7.39 min) whose ionisation is significantly improved by NH_4_F; however, the sensitivity of these steroids was sufficient for their quantitation in serum samples (Figure 2).

**Figure 1.**
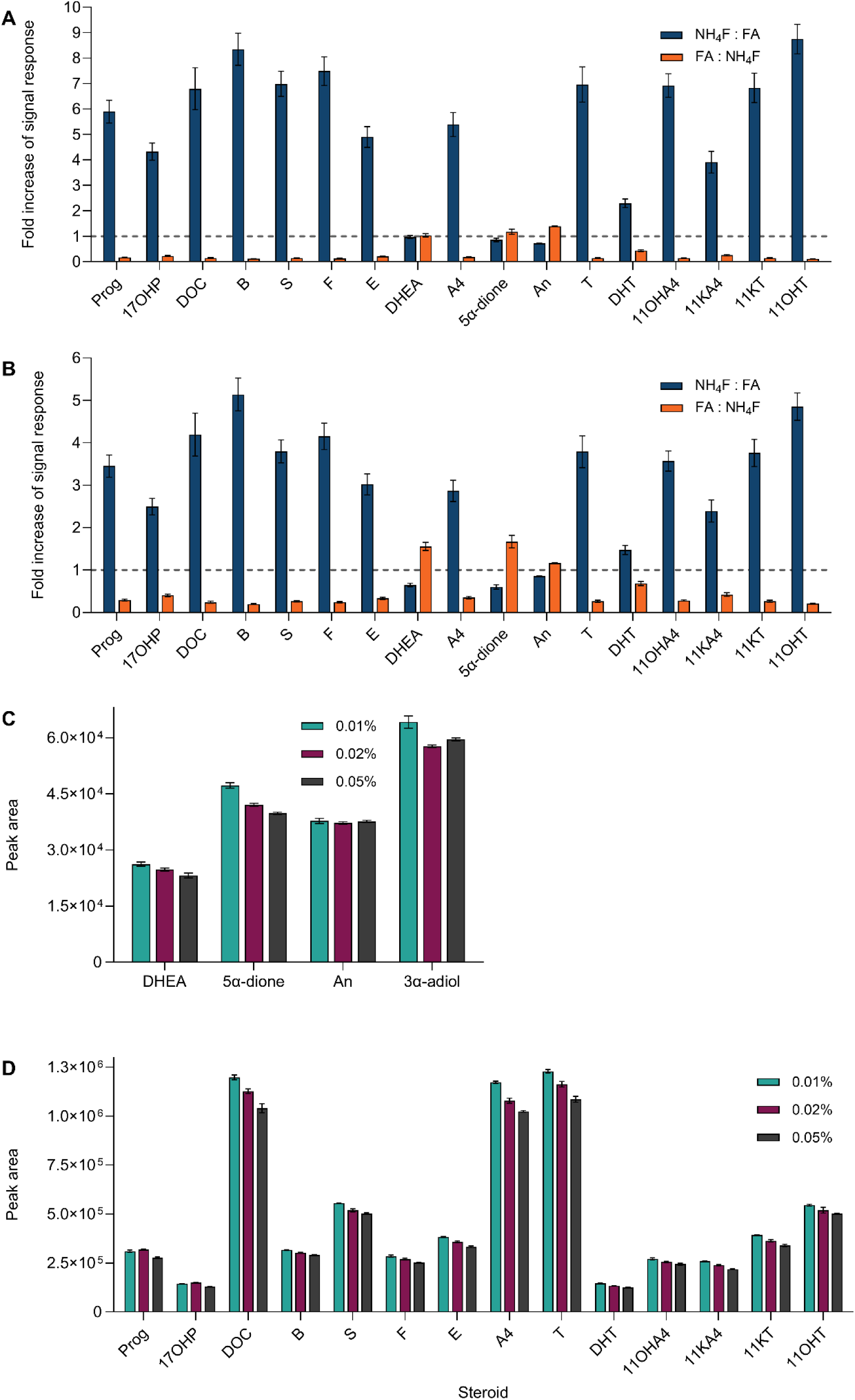
Ionisation efficiency of 2 ng/mL steroid standard solution was evaluated as ratios of peak areas after ammonium fluoride post-column infusion and formic acid in mobile phase (A – without FA in the mobile phase during NH_4_F infusion, B – with FA). Results were also expressed in the opposite ratio to show improved ionisation of A-ring reduced steroids. C, D – different concentrations of FA during timed post-column NH_4_F infusion. Prog – progesterone, 17OHP – 17α-hydroxyprogesterone, DOC – 11-deoxycorticosterone, B – corticosterone, S – 11-deoxycortisol, F – cortisol, E – cortisone, DHEA – dehydroepiandrosterone, A4 – androstenedione, 5α-dione – 5α-androstane-3,17-dione, An – 5α-androsterone, T – testosterone, DHT – 5α-dihydrotestosterone, 3α-adiol – 5α-androstane-3α,17β-diol, 11OHA4 – 11β-hydroxyandrostenedione, 11KA4 – 11-ketoandrostenedione, 11KT – 11-ketotestosterone, 11OHT – 11β-hydroxytestosterone.

**Figure 2.**
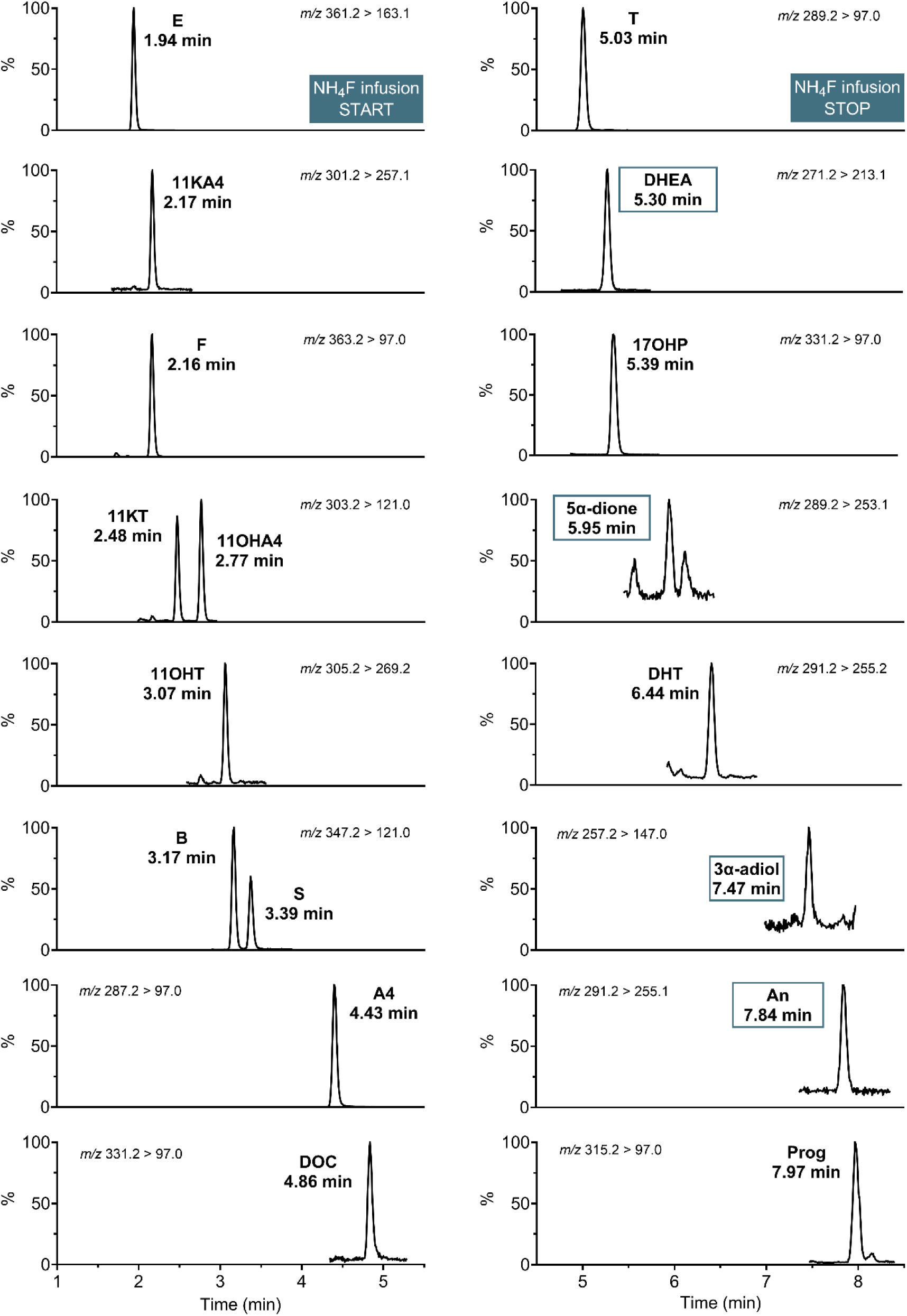
Chromatographic separation of steroids analysed in an extracted serum sample. The post-column ammonium fluoride infusion begins prior to elution of cortisone (E) and ends after elution of testosterone (T; 1.1-5.15 min). The square marks steroids ionising better with formic acid alone. Prog – progesterone, 17OHP – 17α-hydroxyprogesterone, DOC – 11-deoxycorticosterone, B – corticosterone, S – 11-deoxycortisol, F – cortisol, DHEA – dehydroepiandrosterone, A4 – androstenedione, 5α-dione – 5α-androstane-3,17-dione, An – 5α-androsterone, DHT – 5α-dihydrotestosterone, 3α-adiol – 5α-androstane-3α,17β-diol, 11OHA4 – 11β-hydroxyandrostenedione, 11KA4 – 11-ketoandrostenedione, 11KT – 11-ketotestosterone, 11OHT – 11β-hydroxytestosterone.

Concentrations of FA and NH_4_F were optimised to be as low as possible to avoid any deleterious effects in the LC-MS/MS system. Mobile phase A (water) and mobile phase B (methanol) were tested at the following FA concentrations: 0, 0.01, 0.02, and 0.05%. The signal response was generally increasing with decreasing concentration of FA in the mobile phase (Figure 1C, D). Therefore, in the final optimised procedure, we selected mobile phases containing 0.01% FA in combination with 2 mmol/L NH_4_F in 50% methanol/water (v/v) post-column infusion at flow rate of 5 µL/min.

### 3.3 Optimisation of Supported-Liquid Extraction

We subsequently optimised the SLE sample preparation by evaluating different organic solvents and solvent volumes. The solvents tested were MTBE, EtOAC, DCM, and mixtures of MTBE and EtOAc (1:1, 4:1), and EtOAc and hexane (3:1) to assess the effectiveness of different solvent polarities. The extraction efficiency was evaluated by comparing signal responses of a 2 ng/mL spike in steroid depleted serum and a pooled serum sample. Supplementary Figure S2 shows that MTBE and MTBE: EtOAc (4:1) had the lowest extraction efficiency. Extraction efficiencies of the remaining solvents were comparable. DCM exhibited marginally superior recoveries in spiked steroid depleted serum (Supplementary Figure S2A, B); however, DCM extraction efficiency in pooled serum samples did not show any differences in comparison to the remaining solvents (Supplementary Figure S2C, D). Ethyl acetate was selected in the final protocol due to better reproducibility and lower toxicity in comparison to DCM and the solvent mixtures. Different elution volumes were evaluated, i.e., 1 x 1 mL, 2 x 0.5 mL, and 2 x 0.9 mL. However, the 1 mL elution performed similarly to the alternatives as judged by extraction recovery and was therefore selected for the final protocol.

### 3.4 Validation of the UHPLC-MS/MS Method

Specificity was evaluated by comparing retention times of structurally similar compounds. Complete baseline separation was achieved for all tested steroid isobars. Calibrators were prepared in PBS 2% BSA. Calibration curves were analysed over ten different batches and were linear for all analytes (R^2^ > 0.997). The LLOQs and ULOQs were in their target ranges (0.04 to 2.1 nmol/L and 43 to 690 nmol/L, respectively; Table 2). Injection-to-injection carry-over of Cal 9 solvent solution was negligible for both analytes and IS (0.001-9%). Dilution integrity was investigated using 10-fold and 40-fold dilutions of a 2xULOQ QC sample in PBS 2% BSA. The accuracy ranged from 90 to 113% with imprecision ≤3.9%, both of which were within the acceptable range (Supplementary Table S4). Accuracy and imprecision were determined in QC samples prepared in PBS 2% BSA at four concentration levels (LLOQ, low, medium, high). Intra-assay (i.e., 5 replicates within 1 batch) and inter-assay (within 4 days) accuracy was between 82% and 113%. Intra- and inter-assay imprecision was ≤9.9% and ≤19.8%, respectively (Table 3). Accuracy and imprecision were acceptable for all analytes. In addition, imprecision was assessed in spiked serum samples at four concentration levels (approximately equal to the PBS 2% BSA QCs) in triplicate over four separate batches. Intra- and inter-assay imprecision was ≤7.5% and ≤10.5%, respectively (Supplementary Table S5). Absolute recoveries were determined at three concentration levels for each analyte. The mean recoveries were between 83% and 111% (Table 4). IS-normalised matrix effects were between 86% and 111%, which was within the acceptable range (Table 4). Negligible (<10%) matrix effects were observed in haemolysed and lipaemic serum samples. Absolute recovery and matrix effects were comparable between PBS 2% BSA and serum samples. Steroid stability was investigated in serum samples. Steroid concentrations remained within ±15% of their time zero concentrations across three freeze-thaw cycles. Processed samples were stable for up to 72 hours when stored at −20°C, and when kept in the autosampler at 10°C, with bias below ±12% from the target value in both storage conditions. The mean bias for all stability studies is summarised in Table S6.

**Table 2.**
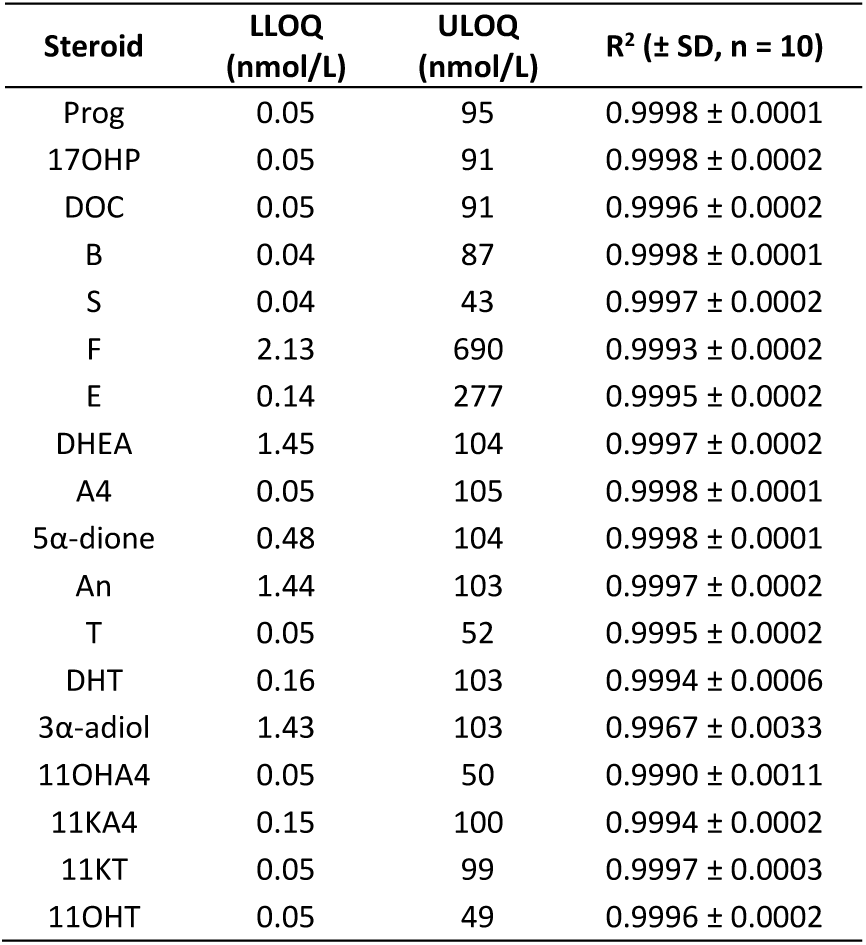
Linear range of analysed steroids (LLOQ – lower limit of quantitation, ULOQ – upper limit of quantitation, SD – standard deviation). Steroid abbreviations as listed in. **Table 1**.

**Table 3.** Intra- and inter-assay accuracy (A) and imprecision (CV). Steroid abbreviations as listed in. **Table 1**.

| Steroid | Target concentration (nmol/L) | Intra-assay (% , n = 5) |  | Inter-assay (% , n = 4) |  | Steroid | Target concentration (nmol/L) | Intra-assay (% , n = 5) |  | Inter-assay (% , n = 4) |  |
| --- | --- | --- | --- | --- | --- | --- | --- | --- | --- | --- | --- |
|  |  | A | CV | A | CV |  |  | A | CV | A | CV |
| Prog | 0.2 | 106 | 2.9 | 106 | 3.3 | 5α-dione | 0.8 | 90 | 6.0 | 90 | 4.8 |
|  | 0.7 | 104 | 3.5 | 104 | 3.6 |  | 2.3 | 93 | 3.8 | 93 | 4.1 |
|  | 38 | 106 | 3.2 | 106 | 7.3 |  | 42 | 95 | 2.9 | 95 | 7.2 |
|  | 76 | 106 | 2.7 | 106 | 7.3 |  | 83 | 95 | 2.3 | 95 | 7.4 |
| 17OHP | 0.2 | 100 | 3.7 | 100 | 4.0 | An | 2.3 | 99 | 3.6 | 99 | 4.7 |
|  | 0.7 | 102 | 3.6 | 102 | 4.1 |  | 6.9 | 101 | 2.9 | 101 | 8.9 |
|  | 36 | 103 | 2.8 | 103 | 8.0 |  | 41 | 102 | 3.2 | 102 | 6.8 |
|  | 73 | 103 | 2.6 | 103 | 8.4 |  | 83 | 101 | 2.7 | 101 | 7.1 |
| DOC | 0.2 | 101 | 2.8 | 101 | 4.0 | T | 0.3 | 98 | 3.2 | 98 | 4.2 |
|  | 0.7 | 101 | 3.2 | 101 | 3.1 |  | 0.8 | 99 | 3.3 | 99 | 3.8 |
|  | 36 | 101 | 2.6 | 101 | 7.3 |  | 21 | 99 | 2.2 | 99 | 7.2 |
|  | 73 | 100 | 2.1 | 100 | 7.2 |  | 42 | 97 | 2.6 | 97 | 7.3 |
| B | 0.2 | 98 | 3.0 | 98 | 5.0 | DHT | 0.3 | 86 | 9.4 | 86 | 13.1 |
|  | 0.6 | 98 | 3.1 | 98 | 3.9 |  | 0.8 | 93 | 3.1 | 93 | 8.0 |
|  | 35 | 100 | 2.5 | 100 | 7.9 |  | 41 | 100 | 3.8 | 100 | 8.6 |
|  | 69 | 99 | 1.9 | 99 | 7.3 |  | 83 | 100 | 3.6 | 100 | 9.2 |
| S | 0.2 | 113 | 2.9 | 113 | 4.7 | 3α-adiol | 2.3 | 82 | 9.9 | 82 | 19.8 |
|  | 0.6 | 111 | 2.9 | 111 | 3.4 |  | 6.8 | 96 | 8.2 | 96 | 11.4 |
|  | 35 | 111 | 2.4 | 111 | 7.0 |  | 41 | 93 | 4.8 | 93 | 6.9 |
|  | 69 | 109 | 1.7 | 109 | 7.0 |  | 82 | 94 | 5.8 | 94 | 8.6 |
| F | 3.4 | 105 | 4.0 | 105 | 4.4 | 11OHA4 | 0.2 | 89 | 5.9 | 89 | 8.8 |
|  | 10 | 108 | 3.5 | 108 | 3.8 |  | 0.7 | 90 | 3.6 | 90 | 3.8 |
|  | 276 | 108 | 2.0 | 108 | 7.5 |  | 40 | 91 | 2.4 | 91 | 7.4 |
|  | 552 | 105 | 2.2 | 105 | 7.5 |  | 79 | 89 | 1.8 | 89 | 7.0 |
| E | 0.7 | 99 | 4.6 | 99 | 5.0 | 11KA4 | 0.2 | 99 | 4.9 | 99 | 5.6 |
|  | 2.1 | 100 | 3.0 | 100 | 3.4 |  | 0.7 | 101 | 3.8 | 101 | 4.1 |
|  | 111 | 102 | 2.2 | 102 | 7.2 |  | 40 | 103 | 2.5 | 103 | 7.5 |
|  | 222 | 100 | 1.9 | 100 | 7.3 |  | 80 | 100 | 1.6 | 100 | 7.2 |
| DHEA | 2.3 | 97 | 5.1 | 97 | 5.5 | 11KT | 0.2 | 104 | 4.6 | 104 | 4.6 |
|  | 6.9 | 100 | 3.1 | 100 | 8.2 |  | 0.7 | 100 | 3.1 | 100 | 3.8 |
|  | 42 | 101 | 2.7 | 101 | 8.1 |  | 40 | 101 | 2.4 | 101 | 7.2 |
|  | 83 | 101 | 2.7 | 101 | 7.1 |  | 79 | 99 | 1.9 | 99 | 7.4 |
| A4 | 0.3 | 101 | 2.7 | 101 | 5.3 | 11OHT | 0.2 | 103 | 7.1 | 103 | 10.0 |
|  | 0.8 | 99 | 2.9 | 99 | 3.4 |  | 0.7 | 108 | 4.2 | 108 | 4.7 |
|  | 42 | 100 | 2.5 | 100 | 7.4 |  | 39 | 110 | 2.6 | 110 | 8.2 |
|  | 84 | 98 | 1.9 | 98 | 7.4 |  | 79 | 107 | 1.7 | 107 | 7.8 |

**Table 4.**
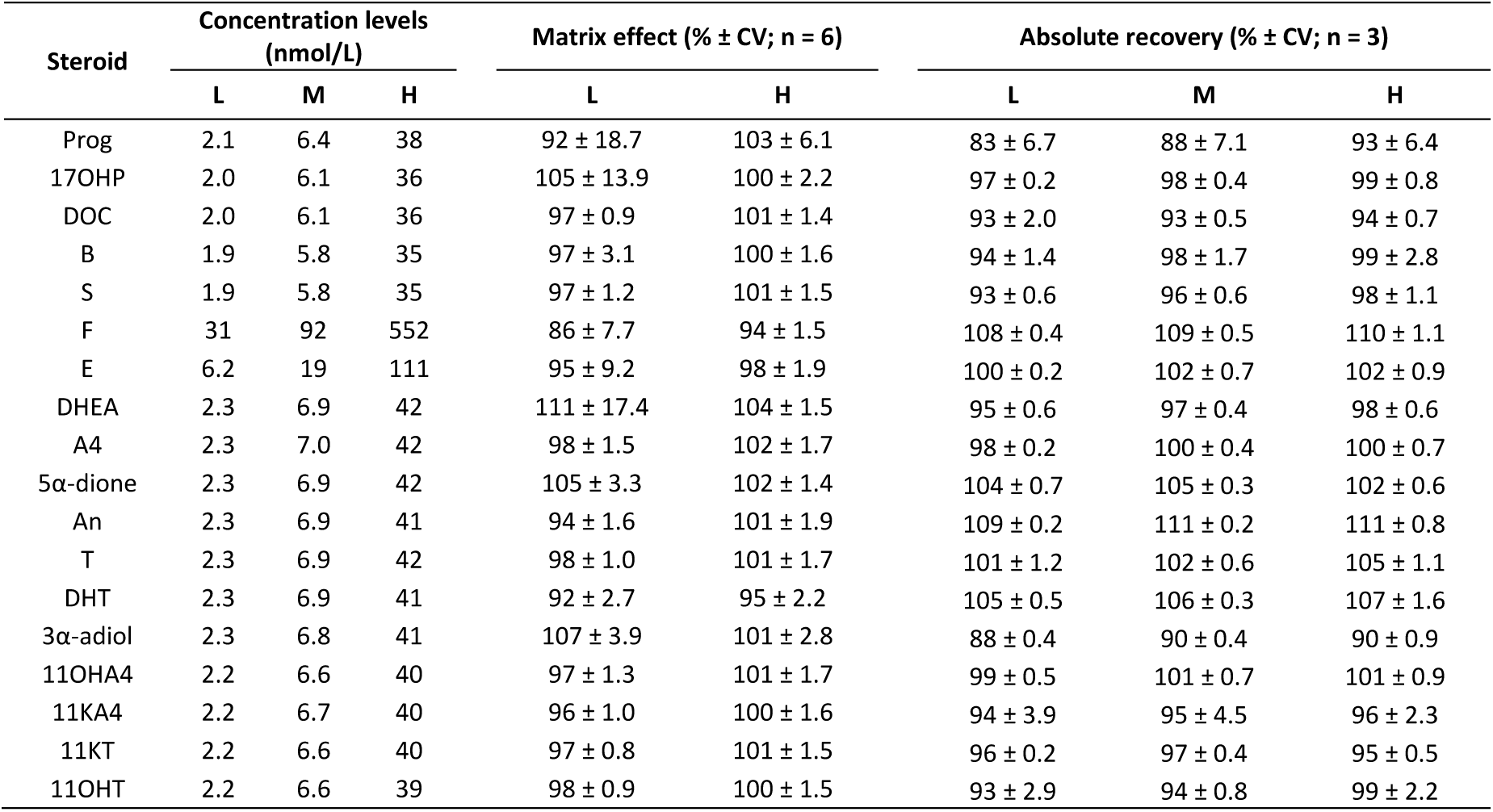
Matrix effect and absolute recovery (L – low, M – medium, H – high, CV – imprecision). Steroid abbreviations as listed in. **Table 1**.

### 3.5 Serum Steroid Profiling in Healthy Individuals

The validated assay was used to quantify serum steroids in healthy, age-matched volunteers of both sexes (55 women, median age 37 (range 22-55) years; 44 men, median age 38 (range 22-60) years). The majority of steroid concentrations were within the linear range, except for 3α-adiol, 5α-dione and An that were below LLOQ. Figure 3 shows the distribution of serum steroid concentrations in women and men. In addition, median steroid concentrations, interquartile ranges (25^th^ – 75^th^ percentiles) and detected concentration ranges are summarised in Table 5. Due to the limited sample size, we did not report the reference intervals. The concentration range of quantified steroids in serum was quite wide; median concentrations of the major serum steroids, cortisol and cortisone, were ≈280 nmol/L and ≈60 nmol/L, respectively. In addition, serum corticosterone, DHEA and, in men, testosterone were other abundant steroids (median concentration <20 nmol/L). Conversely, the median concentrations of the remaining steroids were below 4 nmol/L (Table 5). The concentrations of A4, DHEA, DOC and Prog were significantly higher in women, while testosterone and DHT were significantly higher in men (Figure 3).

**Figure 3.**
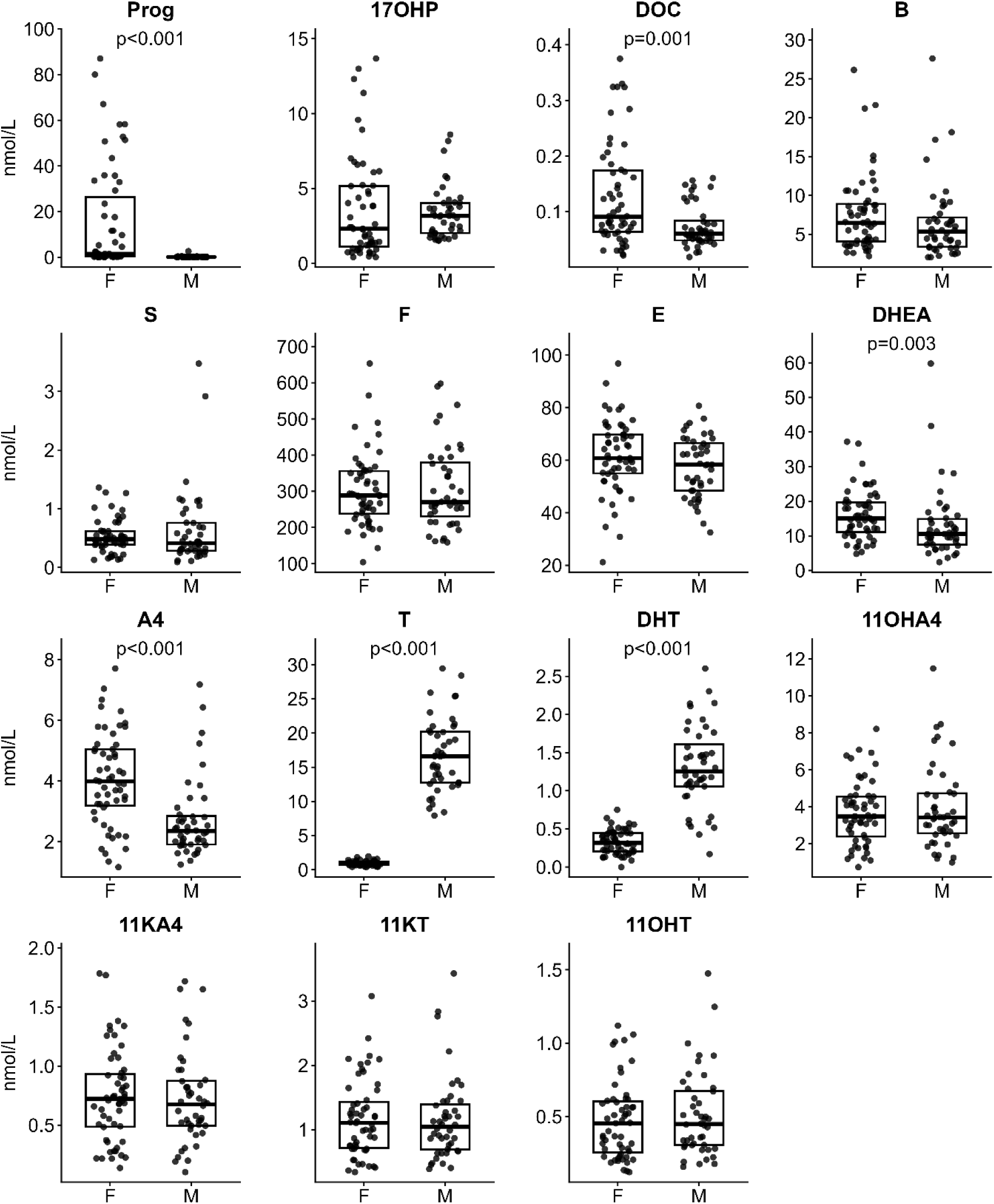
Distribution of steroid concentrations in serum from female (F; n = 55) and male (M; n= 44) healthy volunteers (boxplots show median, the lower and upper quartile). Mann-Whitney U test was used to compare steroid concentrations between men and women. Only p-values of statistically significant results are shown, i.e., p < 0.05. Prog – progesterone, 17OHP – 17α-hydroxyprogesterone, DOC – 11-deoxycorticosterone, B – corticosterone, S – 11-deoxycortisol, F – cortisol, E – cortisone, DHEA – dehydroepiandrosterone, A4 – androstenedione, T – testosterone, DHT – 5α-dihydrotestosterone, 11OHA4 – 11β-hydroxyandrostenedione, 11KA4 – 11-ketoandrostenedione, 11KT – 11-ketotestosterone, 11OHT – 11β-hydroxytestosterone.

**Table 5.** Age (years) and serum steroid concentrations (nmol/L) of healthy individuals (IQR – interquartile range, <LLOQ – below lower limit of quantitation). Steroid abbreviations as listed in. **Table 1**.

| Variable | Women |  |  | Men |  |  |
| --- | --- | --- | --- | --- | --- | --- |
|  | Median | IQR | Range (min - max) | Median | IQR | Range (min - max) |
| Age | 37 | 33 - 41 | 22 - 55 | 38 | 31 - 45 | 22 - 60 |
| Prog | 1.52 | 0.37 - 26.4 | 0.05 - 87.1 | 0.12 | 0.10 - 0.22 | 0.06 - 2.67 |
| 17OHP | 2.31 | 1.11 - 5.17 | 0.41 - 13.67 | 3.18 | 2.03 - 4.03 | 1.52 - 8.60 |
| DOC | 0.11 | 0.08 - 0.19 | <LLOQ - 0.38 | 0.07 | 0.05 - 0.11 | <LLOQ - 0.16 |
| B | 6.47 | 4.07 - 8.89 | 2.19 - 26.1 | 5.34 | 3.38 - 7.16 | 2.05 - 27.6 |
| S | 0.48 | 0.39 - 0.62 | 0.13 - 1.36 | 0.41 | 0.28 - 0.76 | 0.09 - 3.48 |
| F | 288.4 | 237.9 - 356.0 | 104.3 - 653.2 | 269.9 | 231.2 - 380.1 | 159.1 - 597.7 |
| E | 60.8 | 55.0 - 69.8 | 21.2 - 96.9 | 58.4 | 48.4 - 66.6 | 32.5 - 80.8 |
| DHEA | 15.1 | 11.1 - 19.7 | 4.8 - 37.2 | 10.6 | 7.50 - 14.9 | 2.41 - 59.8 |
| A4 | 3.99 | 3.19 - 5.05 | 1.17 - 7.71 | 2.36 | 1.91 - 2.84 | 1.25 - 7.18 |
| T | 0.89 | 0.73 - 1.17 | 0.39 - 1.97 | 16.6 | 12.8 - 20.2 | 7.93 - 29.5 |
| DHT | 0.35 | 0.24 - 0.45 | <LLOQ - 0.75 | 1.25 | 1.05 - 1.61 | 0.17 - 2.60 |
| 11OHA4 | 3.47 | 2.39 - 4.55 | 0.73 - 8.21 | 3.41 | 2.56 - 4.72 | 0.98 - 11.48 |
| 11KA4 | 0.73 | 0.49 - 0.94 | <LLOQ - 1.78 | 0.69 | 0.50 - 0.88 | <LLOQ - 1.72 |
| 11KT | 1.10 | 0.72 - 1.43 | 0.34 - 3.08 | 1.04 | 0.69 - 1.39 | 0.39 - 3.43 |
| 11OHT | 0.45 | 0.25 - 0.60 | 0.12 - 1.12 | 0.45 | 0.30 - 0.67 | 0.16 - 1.47 |

## 4 Discussion

Many serum steroids occur at low concentrations and exhibit poor ionisation efficiency. Therefore, we developed and validated a robust UHPLC-MS/MS assay for analysis of 18 steroids from a low volume of serum with a focus on enhancing ionisation by investigating the impacts of additives and variations in sample preparation. Our method included a comprehensive panel of classic and 11-oxygenated androgens to enable androgen profiling in women with PMOS, as well as in individuals with disorders characterised by androgen excess or dysregulation, such as congenital adrenal hyperplasia [21,22]. In addition, we included selected glucocorticoids, and precursor steroids that are routinely measured in clinical practice, and which can provide a comprehensive fingerprint of steroidogenesis.

We based the development of this assay on our previously published method [23] that introduced the post-column infusion of NH_4_F to prevent column degradation and gradual increase in system pressure that is observed with the addition of NH_4_F in the mobile phase [24]. Ammonium fluoride has previously been shown to significantly improve ionisation efficiency of steroids in both positive and negative mode depending on their molecular structure [11,24–26].

Here, we observed that a post-column infusion of 2 mmol/L NH_4_F significantly improved ionisation efficiency of 3-keto-Δ4 steroids and DHT in comparison to 0.01% FA in the mobile phase, while remaining A-ring reduced steroids (DHEA, An, 3α-adiol, 5α-dione) exhibited higher ionisation efficiency with FA. This latter observation is consistent with a previous study by Galmiche et al. [25] who reported decreased signal for A-ring reduced steroids, as well as for bile acids, steroid sulphates and glucuronides. We therefore implemented a timed post-column infusion stopping the NH_4_F infusion before the elution of A-ring reduced steroids. This approach allowed us to enhance the ionisation of 3-keto-Δ4 steroids, which eluted during the first part of the run, while preventing the suppression of the later eluting poorly ionising A-ring reduced steroids. Gazárková et al. [11] previously suggested that NH_4_F could be useful for a selective tuning of the 3-keto-Δ4 steroids response. Our study is the first to establish a timed post-column infusion for optimised detection across a large range of steroids.

We also optimised sample preparation for a small serum volume (100 µL) by 96-well plate SLE without subsequent derivatisation thereby facilitating high sample throughput. Small sample volumes are desirable in both the clinical and research settings where sample availability is limited or when multiple analyses are required, while also contributing to reduced analysis cost. Importantly, the requirement for smaller volumes also minimises patient and donor discomfort, which is especially beneficial in paediatric and elderly populations.

SLE is a two-step process including sample loading into the SLE sorbent and elution using water-immiscible solvents; in this study we used ethyl acetate. This makes SLE simpler and faster than SPE. Only a few previous studies employed SLE for serum steroid analysis; these studies utilised MTBE or DCM for steroid elution [12,27,28]. It should be noted that during sample preparation we avoided heating samples to prevent non-enzymatic conversion of cortisol and cortisone to 11OHA4 and 11KA4, respectively [29].

We subsequently used the validated assay to quantify steroids in serum samples from age-matched female and male healthy individuals. We found significantly higher levels of Prog, DOC, DHEA, and A4 in women, and higher concentrations of T and DHT in men. Steroid concentrations were within the reference intervals of healthy populations previously determined by LC-MS/MS methods [7,30–36]. These studies are summarised in Table 6 to show comparison of study cohorts, exclusion criteria and analysed steroids. In addition, steroids previously reported to exhibit significant sex differences are highlighted in Table 6. It has been well established that circulating levels of Prog, T, and DHT differ significantly between sexes [31]. Progesterone changes during the menstrual cycle and peaks during the luteal phase. In addition to Prog, we also observed higher levels of DOC in women that may also be a menstrual cycle-driven difference. We previously demonstrated higher Prog and 17OHP levels in saliva in the luteal phase as compared to the follicular phase [35]. When recruiting our healthy control cohort, we established that premenopausal women had regular menstrual cycles, but we did not record the specific cycle day on the day of blood sampling. Frederiksen et al. [34] analysed serum steroids of almost 2500 participants aged 0.17 to 77 years, and they reported significantly higher concentration of androgens in men (i.e., T, DHT, DHEA) except for A4. They described higher levels of A4 in women in early adulthood when the concentration of A4 peaked and was followed by gradual decrease to less than half in late adulthood and reaching similar levels in both sexes. In contrast to our findings, Frederiksen et al. [34] found similar concentrations of DOC in men and women throughout the lifetime [34], which was also reported by Anker et al. [31]. In addition, Anker et al. showed similarly to this study higher concentrations of DHEA and A4 in women [31]. Davio et al. [33] also reported higher A4 concentrations in women. By contrast, Schiffer et al. [35] showed no differences in DHEA and A4 concentrations between sexes in serum samples from 290 participants. In addition, Schiffer et al. reported that, unlike classic androgens, levels of 11-oxygenated androgens did not differ between sexes which we also observed in this study. In contrast, Zeidler et al. [36] reported slightly higher levels of 11OHA4 and 11KA4 in males between 20 and 60 years of age. In summary, these studies show that for comprehensive determination of steroid reference intervals, it is necessary to include and consider multiple parameters including sex, age, BMI, menstrual cycle and hormonal contraceptives [32,34–38].

**Table 6.** Previously published studies on serum steroid reference intervals by LC-MS/MS in healthy population (^F, M^ – steroid was reported higher in females and males, respectively). Steroid abbreviations as listed in. **Table 1**.

| Paper | Participants | Cohort | Exclusion criteria | Variables | Steroid panel | Additional information |
| --- | --- | --- | --- | --- | --- | --- |
| Ahamed et al. (2026) [30] | 500 (250 M, 250 F)<br>19-70 years | Blood transfusion department (China) | Endocrine diseases, hormone replacement therapy, oral contraceptives, corticosteroid use, chronic illnesses, pregnancy, excessive smoking and alcohol intake, extreme physical training | age, sex | Prog<sup>F</sup>, T<sup>M</sup>, estradiol<sup>F</sup> | Steroid hormones declined with age (estradiol levels similar between sexes at the ages of 50 to 70). |
| Anker et al. (2022) [31] | >10,000 (≈70% F),<br>0-100 years | Routine laboratory results (Germany) | Serum sample - low volume, haemolysis, icterus or lipaemia | age, sex | 17α-hydroxypregnenolone, Prog<sup>F</sup>, 17OHP, DOC, B, aldosterone, S, F, E, DHEA<sup>F</sup>, A4<sup>F</sup>, T<sup>M</sup>, DHT<sup>M</sup> | No information about endocrine diseases, malignancies, glucocorticoid therapy, oral contraceptives, puberty state or menopause. |
| Bae et al. (2019) [32] | 4678 (2512 M, 2166 F)<br>0.3-79 years | LIFE Child + Adult studies (Germany) | Endocrine diseases, glucocorticoid therapy, hepatitis, pregnancy, BMI ≥ 33 | age, sex, puberty | Prog<sup>M</sup>, 17OHP, aldosterone, F, E, DHEA sulphate<sup>F</sup>, A4<sup>F</sup>, T, estradiol | Sex differences were reported for the first year of life (mini-puberty). |
| Davio et al. (2020) [33] | 590 (319 M, 271 F)<br>18-97 years | Routine laboratory results (USA) | Systemic steroids therapy, menstrual cycle irregularities, active malignancies or hypogonadism | age, sex, BMI | F<sup>F</sup>, E, A4<sup>F</sup>, T<sup>M</sup>, 11OHA4<sup>M</sup>, 11KA4<sup>M</sup>, 11KT<sup>M</sup>, 11OHT | 11KA4 and 11KT declined with aging only in men. |
| Fanelli et al. (2011) [7] | 416 (217 M, 199 F)<br>19-89 years | Routine laboratory results (Italy) | Endocrine, hepatic, renal, tumoral, autoimmune, cardiovascular, hematologic, neurologic or psychiatric diseases, sleep disorders | age, sex, menstrual cycle | Prog, 17OHP, DOC, B, S, F, DHEA, A4, T | Age-dependent decrease of androgens was shown in both sexes. |
| Frederiksen et al. (2024) [34] | 2458 (1282 M, 1176 F)<br>0.17-77 years | Five cohort studies (Denmark) | Not reported | age, sex | 17-hydroxypregnenolone, Prog<sup>F</sup>, 17OHP<sup>M</sup>, DOC, B, aldosterone, 21-deoxycortisol, S, F<sup>F</sup>, E, DHEA<sup>M</sup>, DHEA sulphate<sup>M</sup>, A4, T<sup>M</sup>, DHT<sup>M</sup>, estrone sulfate | Cortisol was higher in women in late puberty and early adulthood. |
| Schiffer et al. (2023) [35] | 290 (125 M, 165 F)<br>21-95 years | Recruitment through advertisement | Diseases affecting steroid biosynthesis or metabolism, medication that interferes with steroid biosynthesis or metabolism except for hormonal contraceptives | age, sex, BMI, menstrual cycle | DHEA, DHEA sulphate, A4, T<sup>M</sup>, DHT<sup>M</sup>, 11OHA4, 11KA4, 11KT | Classic but not 11-oxygenated androgens declined with age. |
| Zeidler et al. (2026) [36] | 2505 (1280 M, 1225 F)<br>0.25-80 years | LIFE Child + Adult studies (Germany) | Hepatic, renal, liver, and endocrine diseases, current inflammation or cancer, oral contraceptives, glucocorticoid treatment, 18.5 ≤ BMI ≥ 30 | age, sex, puberty | 11OHA4<sup>M</sup>, 11KA4<sup>M</sup>, 11KT, 11OHT | Slightly higher levels of 11OHA4 and 11KA4 were found in males between 20 and 60 years of age. |

## 5 Conclusions

The UHPLC-MS/MS assay for comprehensive steroid profiling was optimised and validated for accurate quantification of 18 steroids in a sample volume of 100 μL. Sensitive detection of steroids was achieved by combining timed post-column infusion of NH_4_F to enhance ionisation of 3-keto-Δ4 steroids and addition of FA in mobile phase to enhance ionisation of A-ring reduced steroids. The validated assay was applied to quantify serum steroids from both female and male healthy adult individuals; given its use of small sample volumes, the assay is amenable for analysis of paediatric cohorts.

## Acknowledgments

This research was supported by the Medical Research Council (MC_UP_1605/15) and the Wellcome Trust (209492/Z/17/Z).

