## Supplementary Material for "Optimisation and Validation of a UHPLC-MS/MS Assay Using Timed Post-Column Ammonium Fluoride Infusion for Sensitive Serum Steroids Quantitation"

***Table S1*** *Reference steroid standards and internal standards used in the assay († – certified reference material, § – powder, ‡ – solution).*

| Steroid name | Abbreviation | Formula | Supplier | CAS number |
|---|---|---|---|---|
| Progesterone | Prog | C21 H30 O2 | Cerilliant†, Round Rock, TX, USA | 57-83-0 |
| 17α-Hydroxyprogesterone | 17OHP | C21 H30 O3 | Cerilliant† / Toronto Research Chemicals, Inc. §, Vaughan, ON, Canada | 68-96-2 |
| 11-Deoxycorticosterone | DOC | C21 H30 O3 | Cerilliant† | 64-85-7 |
| Corticosterone | B | C21 H30 O4 | Cerilliant† | 50-22-6 |
| 11-Deoxycortisol | S | C21 H30 O4 | Cerilliant† / Lipomed†, Arlesheim, Switzerland | 152-58-9 |
| Cortisol | F | C21 H30 O5 | Cerilliant† / Lipomed† | 50-23-7 |
| Cortisone | E | C21 H28 O5 | Cerilliant† | 53-06-5 |
| Dehydroepiandrosterone | DHEA | C19 H28 O2 | Cerilliant† | 53-43-0 |
| Androstenedione | A4 | C19 H26 O2 | Cerilliant† / Lipomed† | 63-05-8 |
| 5α-Androstane-3,17-dione | 5α-dione | C19 H28 O2 | Toronto Research Chemicals§ / Steraloids§, Newport, RI, USA | 846-46-8 |
| 5α-Androsterone | An | C19 H30 O2 | Steraloids§ | 53-41-8 |
| Testosterone | T | C19 H28 O2 | Cerilliant† | 58-22-0 |
| 5α-Dihydrotestosterone | DHT | C19 H30 O2 | Cerilliant† | 521-18-6 |
| 5α-Androstane-3α,17β-diol | 3α-adiol | C19 H32 O2 | Merck§, Gillingham, UK | 1852-53-5 |
| 11β-Hydroxyandrostenedione | 11OHA4 | C19 H26 O3 | Toronto Research Chemicals§ / Merck§ | 382-44-5 |
| 11-Ketoandrostenedione | 11KA4 | C19 H24 O3 | Merck§ | 382-45-6 |
| 11-Ketotestosterone | 11KT | C19 H26 O3 | Cayman Chemical†, Ann Arbor, MI, USA | 564-35-2 |
| 11β-Hydroxytestosterone | 11OHT | C19 H28 O3 | Steraloids§ | 1816-85-9 |
| **Internal standards** | | | | |
| Progesterone-2,3,4-$^{13}$C3 | Prog-$^{13}$C3 | $^{13}$C3 C18 H30 O2 | Merck‡ | 327048-87-3 |
| 17α-Hydroxyprogesterone-2,3,4-$^{13}$C3 | 17OHP-$^{13}$C3 | $^{13}$C3 C18 H30 O3 | Cerilliant† | 1356154-92-1 |
| 11-Deoxycorticosterone-2,3,4-$^{13}$C3 | DOC-$^{13}$C3 | $^{13}$C3 C18 H30 O3 | Merck‡ | 2482452-16-2 |
| Corticosterone-$^{13}$C4 | B-$^{13}$C4 | C17 $^{13}$C4 H30 O4 | ASI Chemicals§, Cheyney, PA, USA | N/A |
| 11-Deoxycortisol-2,3,4-$^{13}$C3 | S-$^{13}$C3 | $^{13}$C3 C18 H30 O4 | Merck‡ | 2482467-26-3 |
| Cortisol-9,11,12,12-d4 | F-d4 | C21 H26 D4 O5 | Cambridge Isotope Laboratories, Inc.§, Andover, MA, USA | 73565-87-4 |
| Cortisone-$^{13}$C3 | E-$^{13}$C3 | C18 $^{13}$C3 H28 O5 | Cerilliant† | N/A |
| Dehydroepiandrosterone-2,2,3,4,4-d5 | DHEA-d5 | C19 H23 D5 O2 | Cerilliant† | 97453-25-3 |
| Androstene-3,17-dione-2,3,4-$^{13}$C3 | A4-$^{13}$C3 | $^{13}$C3 C16 H26 O2 | Cerilliant† | 327048-86-2 |
| 5α-Androstane-3, 17-dione-2,3,4-$^{13}$C3 | 5α-dione-$^{13}$C3 | $^{13}$C3 C16 H28 O2 | Cambridge Isotope Laboratories§ | 2483735-09-5 |
| Androsterone-d4 | An-d4 | C19 D4 H26 O2 | IsoSciences§, Ambler, PA, USA | 89685-10-9 |
| Testosterone-2,3,4-$^{13}$C3 | T-$^{13}$C3 | $^{13}$C3 C16 H28 O2 | Cerilliant† | 327048-83-9 |
| 5α-Dihydrotestosterone- 16,16,17-d3 | DHT-d3 | C19 H27 O2 D3 | Cerilliant† | 79037-34-6 |
| 5α-Androstane-3α,17β-diol-16,16,17-d3 | 3α-adiol-d3 | C19 2H3 H29 O2 | Toronto Research Chemicals§ | 79037-33-5 |
| 11-Hydroxyandrostenedione-2,3,4-$^{13}$C3 | 11OHA4-$^{13}$C3 | C16 $^{13}$C3 H26 O3 | ASI Chemicals§ | N/A |
| 11-Ketoandrostenedione-$^{13}$C3 | 11KA4-$^{13}$C3 | $^{13}$C3 C16 H24 O3 | ASI Chemicals§ | N/A |
| 11-Ketotestosterone-2,3,4-$^{13}$C3 | 11KT-$^{13}$C3 | C16 $^{13}$C3 H26 O3 | ASI Chemicals§ | N/A |
| 11β-Hydroxytestosterone-2,3,4-$^{13}$C3 | 11OHT-$^{13}$C3 | C16 $^{13}$C3 H28 O3 | ASI Chemicals§ | N/A |

***Table S2*** *Concentration of calibrators prepared using PBS 2% BSA (ng/mL, nmol/L – in brackets).*

| Level | Cortisol | Cortisone | Remaining steroids[†] |
|---|---|---|---|
| **Cal 0** | 0 | 0 | 0 |
| **Cal 1** | <LLOQ | 0.05 (0.1) | 0.02 (0.04-0.05) |
| **Cal 2** | 0.77 (2) | 0.15 (0.4) | 0.05 (0.1-0.2)[§] |
| **Cal 3** | 2.32 (6) | 0.46 (1.3) | 0.14 (0.4-0.5)[‡] |
| **Cal 4** | 6.94 (19) | 1.39 (4) | 0.42 (1.2-1.5)[*] |
| **Cal 5** | 20.83 (57) | 4.17 (12) | 1.25 (3.6-4.4) |
| **Cal 6** | 62.50 (172) | 12.50 (35) | 3.75 (11-13) |
| **Cal 7** | 125 (345) | 25 (69) | 7.50 (22-26) |
| **Cal 8** | 250 (690) | 50 (139) | 15 (43-52) |
| **Cal 9** | 500 (1379) | 100 (277) | 30 (87-105) |

[†] The lowest calibrators were below lower limit of quantitation (<LLOQ) for some steroids and were excluded during sample analyses.

[§] LLOQ for 11-ketoandrostenedione and 5α-dihydrotestosterone.

[‡] LLOQ for 5α-androstane-3,17-dione.

[*] LLOQ for 5α-androstane-3α,17β-diol, 5α-androsterone, and dehydroepiandrosterone.

***Table S3*** *Concentration of quality control (QC) samples prepared using PBS 2% BSA (ng/mL, nmol/L - in brackets).*

| Levels | Cortisol | Cortisone | Remaining steroids |
|---|---|---|---|
| **QC 1** | 1.24 (3.4) | 0.25 (0.7) | 0.07 (0.2-0.3) |
| **QC 2** | 3.70 (10) | 0.74 (2.1) | 0.22 (0.6-0.8) |
| **QC 3** | 11.11 (31) | 2.22 (6.2) | 0.67 (1.9-2.3) |
| **QC 4** | 33.33 (92) | 6.66 (19) | 2 (5.8-7.0) |
| **QC 5** | 100 (276) | 20 (56) | 6 (178-21) |
| **QC 6** | 200 (552) | 40 (111) | 12 (35-42) |
| **QC 7** | 400 (1104) | 80 (222) | 24 (69-84) |

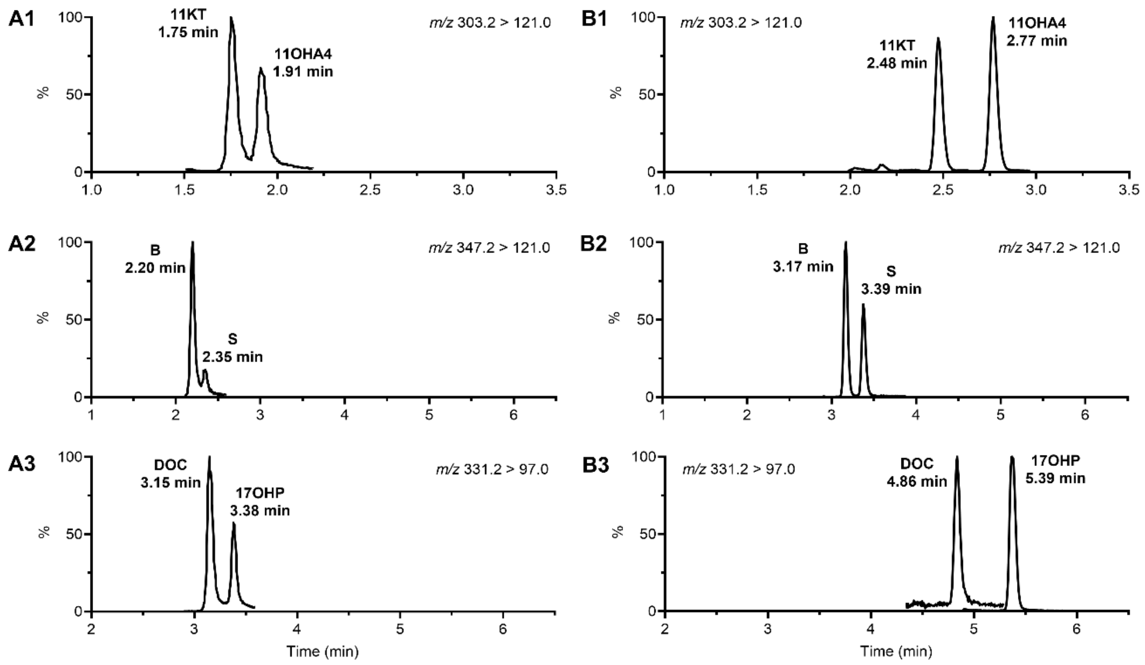


***Figure S1*** *Steroid separation using 50 mm (A1-A3) and 100 mm (B1-B3) Luna Omega Polar C18 columns. Steroid abbreviations are explained in Table S1.*

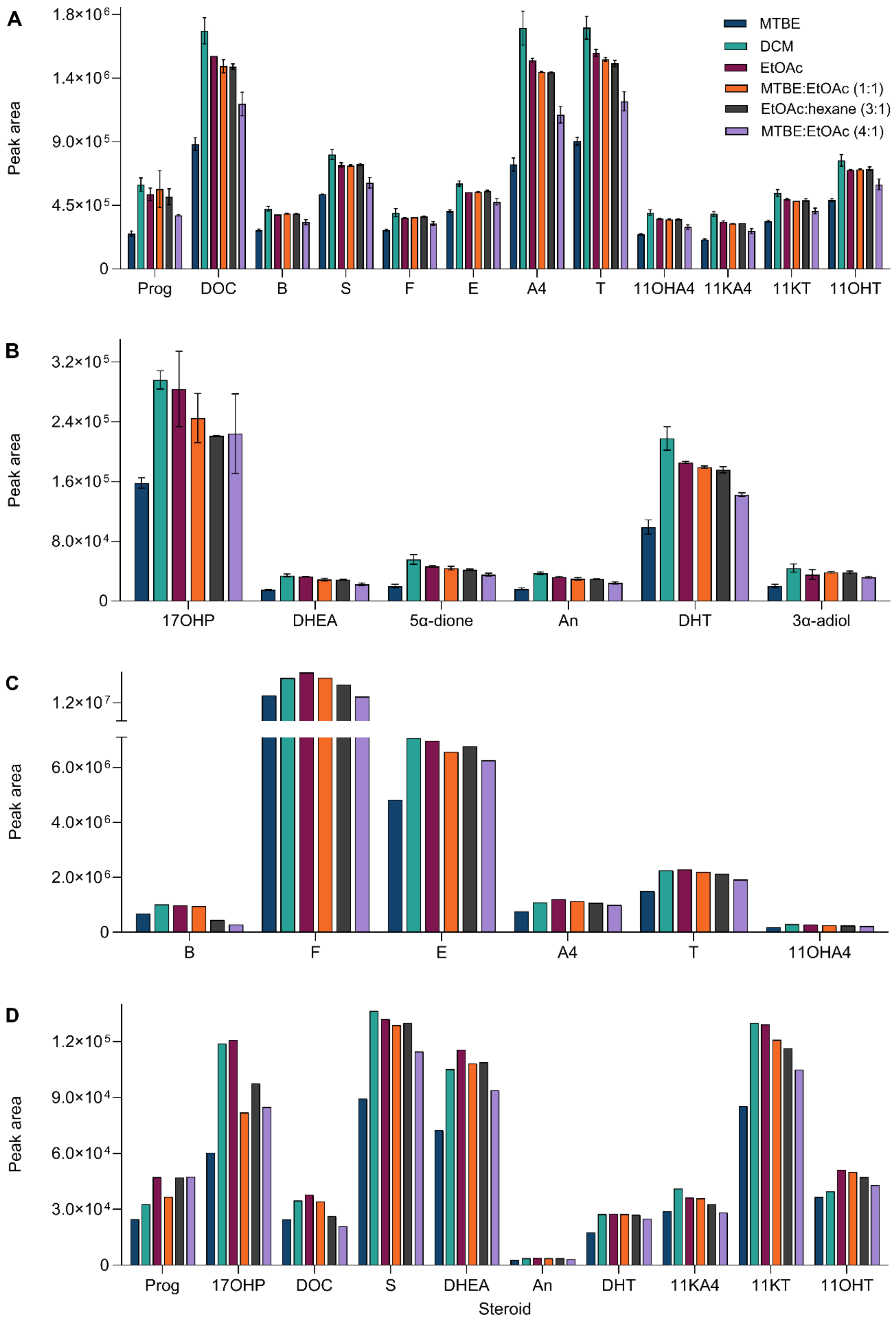


***Figure S2*** *Solvent extraction efficiency. Steroid peak areas in 2 ng/mL spiked steroid depleted serum analysed in duplicate for each elution solvent (A, B) and a pool serum sample (C, D) after sample preparation by supported liquid extraction. Steroid abbreviations are explained in Table S1.*

***Table S4*** *Dilution integrity of 2xULOQ QC. Steroid abbreviations are explained in Table S1.*

| Steroid | Nominal concentration 2xULOQ (nmol/L) | 10-fold dilution | | 40-fold dilution | |
|---|---|---|---|---|---|
| | | Accuracy (%) | Imprecision (%) | Accuracy (%) | Imprecision (%) |
| Prog | 191 | 110 | 0.5 | 108 | 2.5 |
| 17OHP | 182 | 106 | 0.8 | 105 | 1.4 |
| DOC | 182 | 104 | 0.7 | 103 | 2.1 |
| B | 173 | 101 | 0.6 | 101 | 1.2 |
| S | 173 | 113 | 0.8 | 113 | 1.8 |
| F | 2759 | 111 | 1.1 | 109 | 1.8 |
| E | 555 | 103 | 0.5 | 103 | 2.0 |
| DHEA | 208 | 101 | 2.6 | 101 | 2.5 |
| A4 | 209 | 100 | 0.7 | 100 | 1.8 |
| 5α-dione | 208 | 94 | 1.0 | 95 | 2.2 |
| An | 207 | 101 | 1.1 | 103 | 3.1 |
| T | 208 | 101 | 0.5 | 99 | 2.4 |
| DHT | 207 | 98 | 2.9 | 103 | 3.1 |
| 3α-adiol | 205 | 90 | 3.9 | 95 | 3.0 |
| 11OHA4 | 198 | 91 | 1.0 | 90 | 1.8 |
| 11KA4 | 200 | 105 | 0.7 | 104 | 1.6 |
| 11KT | 198 | 101 | 0.3 | 101 | 1.5 |
| 11OHT | 197 | 110 | 1.1 | 110 | 1.9 |

***Table S5*** *Intra- and inter-assay imprecision (CV) of spiked serum samples (n.d. – not detected). Steroid abbreviations are explained in Table S1.*

| **Steroid** | **Target concentration (nmol/L)** | **Intra-assay (n = 3; %)** | **Inter-assay (n = 4; %)** | **Steroid** | **Target concentration (nmol/L)** | **Intra-assay (n = 3; %)** | **Inter-assay (n = 4; %)** |
|---|---|---|---|---|---|---|---|
| Prog | 0 | 0.9 | 5.5 | 5α-dione | 0 | 3.2 | 3.5 |
| | 0.2 | 3.0 | 3.8 | | 0.8 | 4.4 | 4.4 |
| | 0.7 | 1.9 | 2.7 | | 2.3 | 3.4 | 5.2 |
| | 38 | 1.9 | 3.7 | | 42 | 2.1 | 3.9 |
| | 76 | 2.5 | 5.3 | | 83 | 2.4 | 5.4 |
| 17OHP | 0 | 1.7 | 6.0 | An | 0 | 4.2 | 8.3 |
| | 0.2 | 2.5 | 4.4 | | 2.3 | 3.9 | 6.0 |
| | 0.7 | 1.6 | 4.5 | | 6.9 | 3.3 | 6.0 |
| | 36 | 2.1 | 4.1 | | 41 | 3.2 | 5.7 |
| | 73 | 2.4 | 5.2 | | 83 | 3.4 | 4.7 |
| DOC | 0 | 2.5 | 5.3 | T | 0 | 1.1 | 7.3 |
| | 0.2 | 2.0 | 3.5 | | 0.3 | 2.1 | 5.0 |
| | 0.7 | 3.8 | 5.2 | | 0.8 | 1.7 | 6.4 |
| | 36 | 2.4 | 3.8 | | 21 | 1.9 | 5.0 |
| | 73 | 2.2 | 5.2 | | 42 | 2.0 | 4.3 |
| B | 0 | 1.0 | 6.7 | DHT | 0 | 7.5 | 10.5 |
| | 0.2 | 2.2 | 4.9 | | 0.3 | 6.6 | 9.9 |
| | 0.6 | 1.4 | 5.3 | | 0.8 | 5.1 | 7.6 |
| | 35 | 2.1 | 4.3 | | 41 | 4.4 | 4.8 |
| | 69 | 1.9 | 5.0 | | 83 | 4.5 | 5.4 |
| S | 0 | 1.1 | 5.7 | 3α-adiol | 0 | n.d. | n.d. |
| | 0.2 | 1.8 | 3.1 | | 2.3 | 7.3 | 9.9 |
| | 0.6 | 2.8 | 4.2 | | 6.8 | 5.8 | 8.9 |
| | 35 | 2.2 | 4.3 | | 41 | 4.8 | 7.0 |
| | 69 | 2.1 | 5.4 | | 82 | 5.5 | 7.5 |
| F | 0 | 1.1 | 6.4 | 11OHA4 | 0 | 1.3 | 5.8 |
| | 3.4 | 2.0 | 4.6 | | 0.2 | 2.4 | 4.8 |
| | 10 | 1.0 | 5.5 | | 0.7 | 1.7 | 4.4 |
| | 276 | 1.4 | 4.5 | | 40 | 2.1 | 3.6 |
| | 552 | 1.6 | 3.9 | | 79 | 2.0 | 4.7 |
| E | 0 | 0.9 | 5.6 | 11KA4 | 0 | 1.0 | 4.9 |
| | 0.7 | 2.2 | 3.9 | | 0.2 | 2.2 | 3.4 |
| | 2.1 | 1.2 | 4.7 | | 0.7 | 2.3 | 3.4 |
| | 111 | 2.0 | 3.8 | | 40 | 2.9 | 4.2 |
| | 222 | 1.7 | 5.2 | | 80 | 2.5 | 5.4 |
| DHEA | 0 | 1.1 | 5.4 | 11KT | 0 | 1.0 | 5.5 |
| | 2.3 | 2.0 | 5.1 | | 0.2 | 1.9 | 3.5 |
| | 6.9 | 1.6 | 5.0 | | 0.7 | 2.0 | 4.5 |
| | 42 | 2.7 | 3.5 | | 40 | 2.6 | 4.0 |
| | 83 | 2.8 | 5.6 | | 79 | 2.2 | 5.4 |
| A4 | 0 | 1.3 | 6.1 | 11OHT | 0 | 2.3 | 5.7 |
| | 0.3 | 2.2 | 4.1 | | 0.2 | 2.5 | 3.6 |
| | 0.8 | 1.1 | 4.7 | | 0.7 | 3.6 | 7.4 |
| | 42 | 2.1 | 4.0 | | 39 | 2.8 | 4.6 |
| | 84 | 2.3 | 5.3 | | 79 | 2.4 | 6.1 |

***Figure S6*** *Stability of steroids in serum (mean bias ± SD). Freezer and autosampler stability were assessed by reinjecting the plate after storage. Steroid abbreviations are explained in Table S1.*

| **Steroid** | **Freeze-thaw cycle 1** | **Freeze-thaw cycle 2** | **Freeze-thaw cycle 3** | **Freezer (-20°C, 72 hours)** | **Autosampler (+10°C, 72 hours)** |
|---|---|---|---|---|---|
| Prog | 1.8 ± 4.4 | 2.6 ± 5.6 | -2.3 ± 3.8 | -3.2 ± 5.1 | -1.7 ± 3.9 |
| 17OHP | -0.4 ± 6.3 | -1.1 ± 5.9 | -2.3 ± 4.6 | 0.8 ± 0.6 | -0.7 ± 0.7 |
| DOC | 3.5 ± 4.8 | 2.4 ± 4.6 | 0.8 ± 6.1 | 0.6 ± 1.4 | -0.8 ± 1.8 |
| B | 0.3 ± 3.2 | -1.1 ± 1.9 | -1.7 ± 1.6 | -0.1 ± 0.3 | -0.4 ± 0.3 |
| S | 0.4 ± 3.0 | -2.1 ± 2.3 | -3.1 ± 2.0 | 1.9 ± 0.6 | 2.4 ± 1.3 |
| F | 0.0 ± 3.0 | -2.0 ± 2.1 | -4.6 ± 1.7 | -0.1 ± 0.4 | 0.4 ± 0.6 |
| E | 0.9 ± 2.9 | -1.1 ± 2.0 | -2.8 ± 1.7 | 0.1 ± 0.3 | 0.5 ± 0.3 |
| DHEA | 1.1 ± 5.0 | -3.3 ± 3.0 | -3.4 ± 2.4 | -2.3 ± 1.4 | 1.1 ± 1.1 |
| A4 | -0.7 ± 2.8 | -1.8 ± 1.6 | -2.5 ± 1.4 | -0.4 ± 0.1 | 0.9 ± 0.2 |
| 5α-dione | 4.1 ± 5.6 | 6.3 ± 8.0 | 3.1 ± 5.2 | 11.0 ± 8.8 | 6.8 ± 6.1 |
| An | 13.9 ± 5.0 | 5.6 ± 5.6 | 7.4 ± 7.6 | -3.5 ± 3.2 | -3.3 ± 4.9 |
| T | -0.6 ± 2.7 | -2.2 ± 1.8 | -4.8 ± 1.4 | 0.1 ± 0.3 | 1.2 ± 0.7 |
| DHT | 2.9 ± 3.2 | 2.1 ± 4.9 | -0.8 ± 6.4 | 3.6 ± 2.4 | 2.8 ± 5.0 |
| 3α-adiol | 12.3 ± 5.6 | 6.2 ± 9.8 | 12.1 ± 9.9 | 9.2 ± 4.2 | 3.2 ± 1.8 |
| 11OHA4 | -2.0 ± 2.9 | -1.9 ± 2.0 | -3.1 ± 1.8 | 0.9 ± 0.4 | -1.5 ± 0.5 |
| 11KA4 | 2.1 ± 3.1 | -0.3 ± 3.5 | -1.4 ± 2.6 | 1.0 ± 0.6 | 2.6 ± 0.8 |
| 11KT | 1.6 ± 3.4 | 1.5 ± 1.5 | -1.8 ± 1.9 | 0.8 ± 0.8 | 0.8 ± 0.8 |
| 11OHT | 0.2 ± 5.9 | -5.9 ± 5.6 | -4.0 ± 5.7 | 0.1 ± 0.7 | 1.5 ± 0.8 |
